# SCT promoter methylation is a highly discriminative biomarker for lung and many other cancers

**DOI:** 10.1101/018515

**Authors:** Adwait Sathe, Yu-An Zhang, Xiaotu Ma, Pradipta Ray, Daniela Cadinu, Yi-Wei Wang, Xiao Yao, Xiaoyun Liu, Hao Tang, Yunfei Wang, Ying Huang, Changning Liu, Jin Gu, Martin Akerman, Yifan Mo, Chao Cheng, Zhenyu Xuan, Lei Chen, Guanghua Xiao, Yang Xie, Luc Girard, Hongyang Wang, Stephen Lam, Ignacio I Wistuba, Li Zhang, Adi F Gazdar, Michael Q Zhang

## Abstract

Aberrant DNA methylation has long been implicated in cancers. In this work we present a highly discriminative DNA methylation biomarker for non-small cell lung cancers and fourteen other cancers. Based on 69 NSCLC cell lines and 257 cancer-free lung tissues we identified a CpG island in SCT gene promoter which was verified by qMSP experiment in 15 NSCLC cell lines and 3 immortalized human respiratory epithelium cells. In addition, we found that SCT promoter was methylated in 23 cancer cell lines involving >10 cancer types profiled by ENCODE. We found that SCT promoter is hyper-methylated in primary tumors from TCGA lung cancer cohort. Additionally, we found that SCT promoter is methylated at high frequencies in fifteen malignancies and is not methylated in ∼1000 non-cancerous tissues across >30 organ types. Our study indicates that SCT promoter methylation is a highly discriminative biomarker for lung and many other cancers.

## 1 INTRODUCTION

Methylation of the fifth position of the cytosine nucleotide (5mC) is one of the best studied epigenetic modifications and is associated with development in mammals [1]. Due to the difference of biochemical properties between methylated cytosine and unmethylated cytosine, methylated cytosine is sometimes called the “fifth” base [2]. DNA methylation occurs almost exclusively at CpG dinucleotide sites, although non-CpG methylation was recently found to be abundant in the stem cell genome [3]. Since methylated cytosine may spontaneously deaminate to thymine, CpG sites are underrepresented in the human genome, possibly due to DNA methylation during evolution [4]. However, there are still stretches of DNA sequences with over-represented CpG sites, called CpG islands [5]. CpG islands are in general unmethylated and overlap the promoter regions of 60%-70% of all human genes [5]. Moreover, the methylation status of cytosine nucleotides is somatically inheritable through maintenance methyltransferase (e.g., DNMT1), which mostly targets hemimethylated CpG dyads on the parental strand during DNA replication [6].

Lung cancer is the leading cause of cancer-related mortality worldwide and many patients are usually diagnosed at advanced stages [7]. Five-year survival of lung cancer patients after resection strongly depends on tumor stages (67% for stage I vs 23% for stage IV), indicating importance of early diagnosis [8]. Computed tomography (CT) screening is the most recent state-of-art method for early detection [9]. However, it is subject to high false-positive rates [10]. As a result, confirming the presence of lung cancers by invasive biopsy is still a key step for early detection [10]. Molecular biomarkers offer great promise in early detection of lung cancers which in turn can reduce mortality. For example, DNA methylation have been studied in squamous cell lung cancer [11] and lung adenocarcinoma [12][13][14], as well as multiple malignancies [15]. DNA methylation biomarkers for lung cancers are also studied in sputum [16] and plasma [17] for non-invasive procedures. In this report, we describe our search for a near universal lung cancer methylation marker, utilizing the vast amount of publically available data.

## II. RESULT

### A. Initial screening for hyper-methylated biomarkers in non-small cell lung cancer (NSCLC) cell lines

In order to discover DNA methylation markers for non-small cell lung cancers, we obtained HumanMethylation450 (450k) methylation data for 69 cell lines (Table S1) derived from tumors of non-small cell lung cancer patients and 257 primary cancer free lung tissues (Table S2). We applied *t*-test on *M*- values for a restricted search on 390,920 CpG sites detectable in >95% cell lines with no CpG sites within 10 base pairs of known SNPs. We considered probes with FDR level 1e-68 and effect size (i.e., differences of average M-values between cancer and non-malignant/normal samples) to be higher than +4.5 (a positive marker for tumors), and have at least two probes that are consistently hyper-methylated in NSCLC cell lines (Fig.S1-2). The top 2 CGIs were chr11:626728-628037 overlapping with the promoter of secretin gene (SCT; Fig. S1-2, Fig. 1(a), Table S3) and chr2:63283936-63284147 overlapping with the 3’UTR of OTX1 (Table S3). Here we present results on probe cg00249511 in the SCT promoter region in main figures and that of probe cg25774643 can be found in supplementary materials.

**Figure 1.**
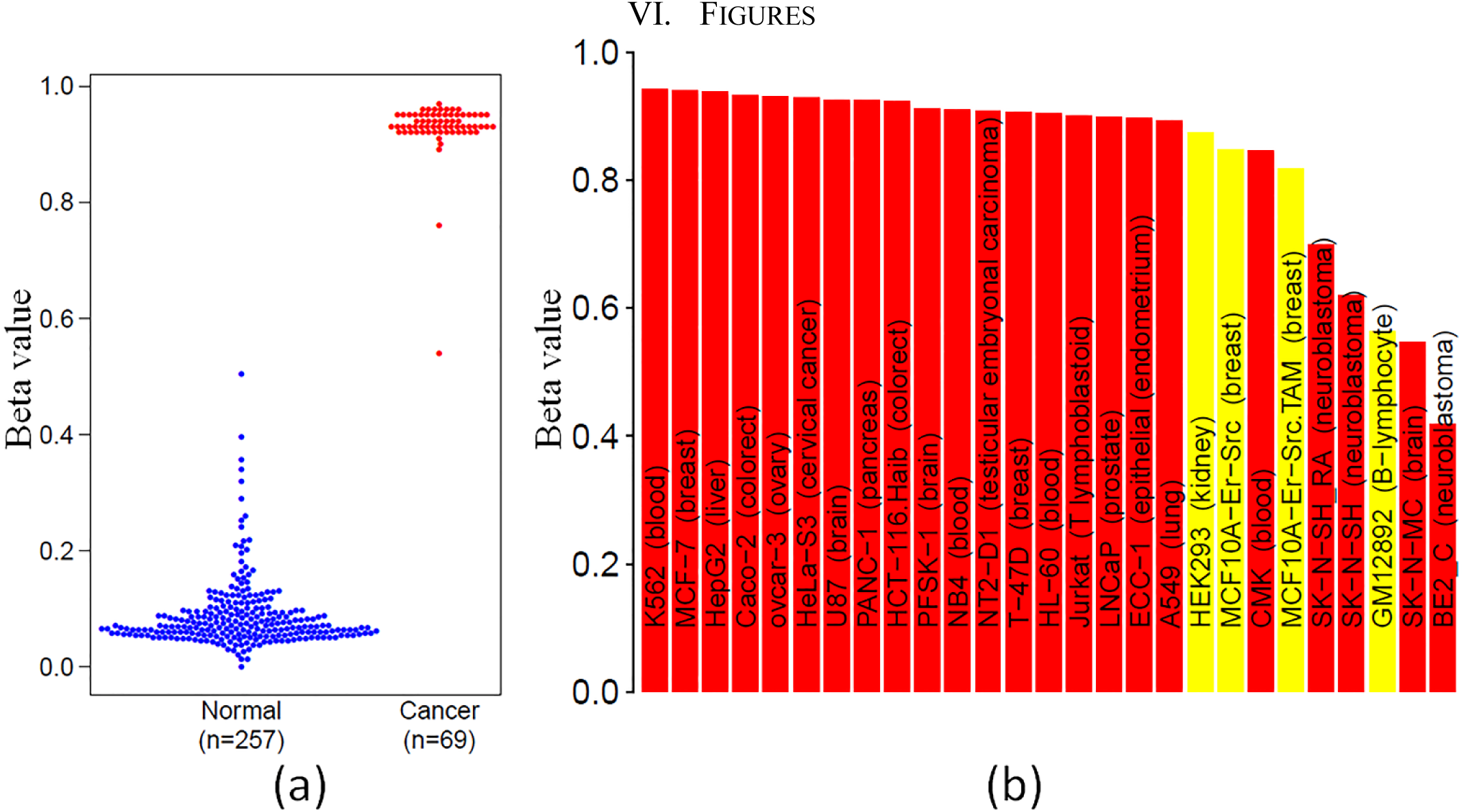
SCT promoter (cg00249511) methylation as a biomarker in (a) Sixty nine NSCLC cell lines (red) and 257 normal lung tissues (blue). Shown in y-axis is the methylation level (β value). (b) SCT methylation in cell lines profiled by ENCODE, including twenty three cancer cell lines (red shade) and four nonmalignant cell lines immortalized by oncogenes or oncogenic viruses agents, such as Epstein-Barr Virus, Adenovirus and ER-Src, which are shaded yellow. Cell line names and tissue sources are indicated.

To check SCT methylation in other cancers we obtained DNA methylation data in cell lines profiled by ENCODE (Table S1), of which 23 cell lines were cancer cell lines. As can be seen in Fig. 1(b) and Fig. S3-4, SCT promoter was methylated in all cancer cell lines, though its methylation level in cell lines BE2 _C and SK _N _MC appears to be reduced. This data indicates that SCT promoter might be a highly discriminative DNA methylation marker for lung and many other cancers.

### B. Validation of SCT promoter methylation by qMSP

To validate the above findings, we developed a SCT-specific qMSP assay (Materials/Methods) by targeting the SCT promoter region and examined 15 NSCLC cancer cell lines and 3 immortalized human respiratory epithelial cell lines, whose SCT methylation status were also confirmed by 450K methylation array (Table S4). As can be seen in Fig. 2 and Fig. S5, the data from both methods consistently indicated SCT promoter being methylated in lung cancer cell lines and none or minimal in those non-malignant lung cell lines. In addition, the SCT qMSP analysis of normal WBC cells showed none or minimal level of SCT methylation, which were consistent with SCT promoter bisulfite sequencing analysis (Zhang et al. manuscript in preparation).

**Figure 2.**
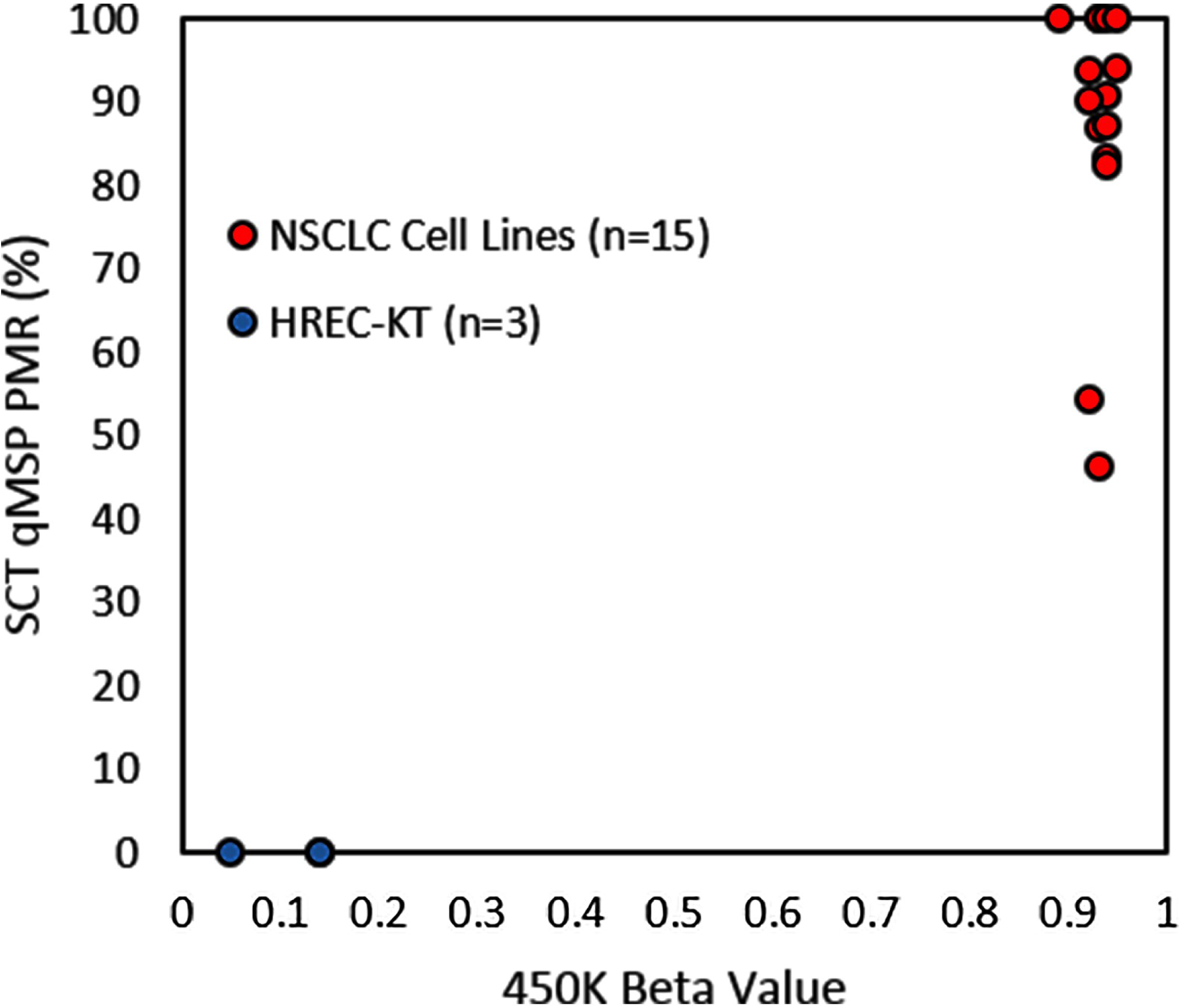
Determination of SCT methylation in genomic DNAs of 15 NSCLC cancer cell lines and 3 immortalized human respiratory epithelial cell lines (HREC-KT) by SCT qMSP assay (y-axis) and 450K methylation array (x-axis; probe cg00249511). PMR: percent of methylated reference in respect to fully methylated DNA controls.

### C. SCT promoter methylation in primary tumors and non-tumor samples

We next checked SCT promoter methylation in 30 cancer types and corresponding non-malignant samples (Table S5) profiled by TCGA [18]. As can be seen in Fig. 3 and Fig. S6 top panel, The SCT promoter is methylated (mean β>0.5) in a wide range of cancers including ovarian, uterine/uterine corpus, cervical, lung, liver, bladder, esophageal, low grade glioma, prostate, head/neck, breast, kidney and stomach, which involves 67.23% (5246/7803) patients in our analysis. Interestingly, SCT promoter is also methylated, though at a reduced level and frequency, in cancers including diffuse large B-cell lymphoma, glioblastoma multiforme, mesothelioma, skin, sarcoma, pancreas, and kidney, which involves 18.65% (1455/7803) patients in our analysis. Finally, SCT promoter does not seem to be methylated in cancers including colorectal, AML, testicular germ cell tumors, adrenocortical, rectum, kidney chromophobe, uveal melanoma and thymoma, which involves 14.12% (1102/7803) patients in our analysis. The descriptive statistics of SCT promoter methylation in all cancer types were summarized in Table 1.

**Figure 3.**
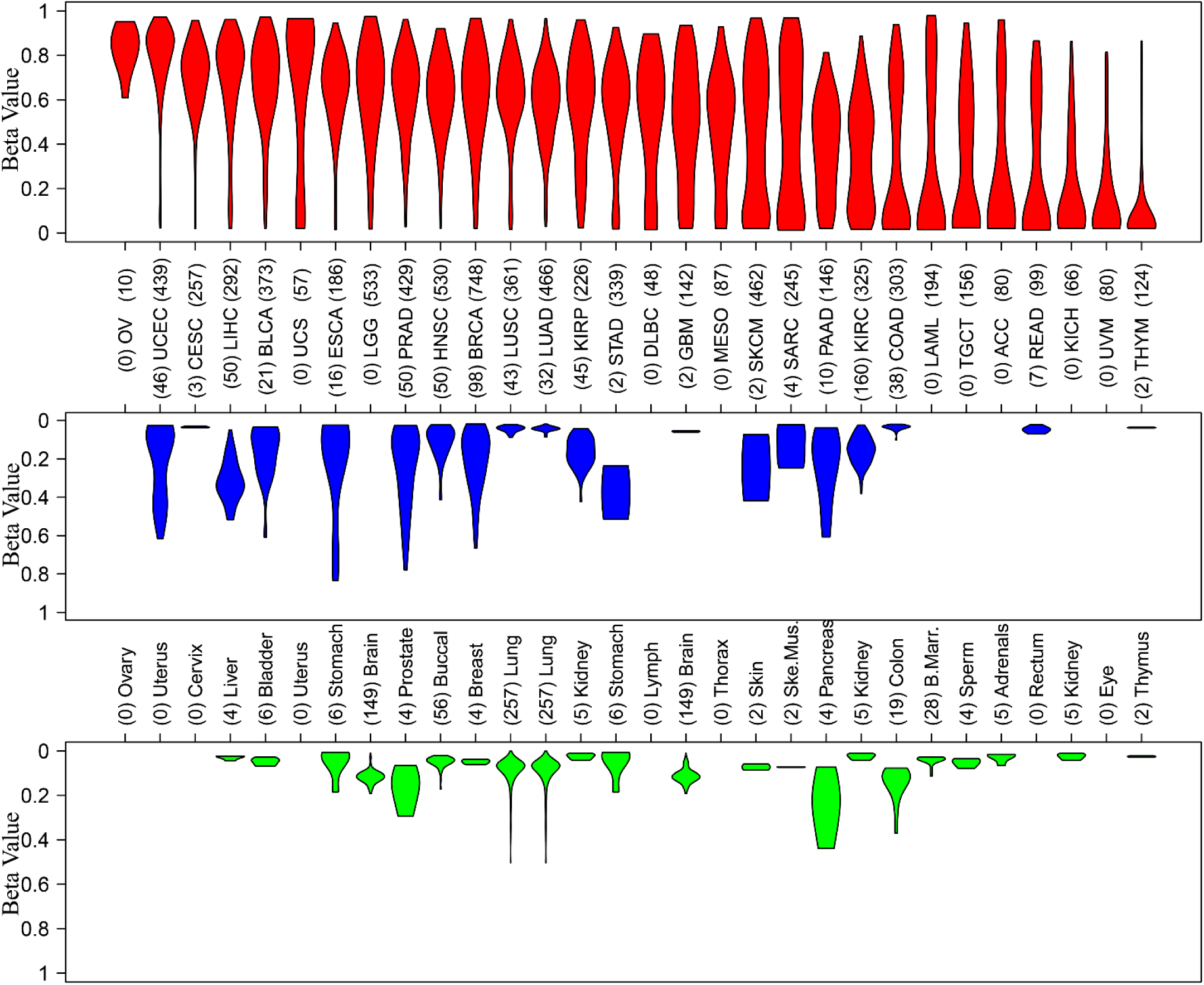
SCT promoter (cg00249511) methylation in TCGA primary tumors (red; top panel), TCGA non-malignant tissues (blue; middle panel) and cancer-free tissues (green; bottom panel). Shown in y-axis is the methylation level (Beta value).

**Table 1.** Summary statistics for SCT promoter methylation in tumor, nonmalignant tissue, and cancer free samples.

| Cancer Type | Sample Sizes | | | cg00249511 ($\beta$) | | | | | | | cg25774643 ($\beta$) | | | | | | |
| --- | --- | --- | --- | --- | --- | --- | --- | --- | --- | --- | --- | --- | --- | --- | --- | --- | --- |
| | Normal | Nonmalig | Tumor | Normal | | Nonmalig | | Tumor | | Δmean (T-N) | Normal | | Normal | | Tumor | | Δmean (T-N) |
|  | (n) | (n) | (n) | Mean | Std | Mean | Std | Mean | Std |  | Mean | Std | Mean | Std | Mean | Std |  |
| OV | 0 | 0 | 10 |  |  |  |  | 0.83 | 0.10 |  |  |  |  |  | 0.86 | 0.10 |  |
| UCEC | 0 | 46 | 439 |  |  | 0.22 | 0.19 | 0.78 | 0.18 | 0.56 |  |  | 0.33 | 0.24 | 0.81 | 0.17 | 0.48 |
| CESC | 0 | 3 | 257 |  |  | 0.03 |  | 0.70 | 0.16 | 0.66 |  |  | 0.08 |  | 0.75 | 0.14 | 0.68 |
| LIHC | 4 | 50 | 292 | 0.03 | 0.01 | 0.31 | 0.11 | 0.68 | 0.22 | 0.37 | 0.02 | 0.01 | 0.65 | 0.09 | 0.76 | 0.18 | 0.10 |
| BLCA | 6 | 21 | 373 | 0.05 | 0.02 | 0.16 | 0.14 | 0.67 | 0.20 | 0.51 | 0.16 | 0.09 | 0.29 | 0.19 | 0.73 | 0.19 | 0.44 |
| UCS | 0 | 0 | 57 |  |  |  |  | 0.67 | 0.34 |  |  |  |  |  | 0.71 | 0.33 |  |
| ESCA | 6 | 16 | 186 | 0.05 | 0.07 | 0.22 | 0.26 | 0.64 | 0.18 | 0.42 | 0.06 | 0.11 | 0.30 | 0.27 | 0.71 | 0.16 | 0.41 |
| LGG | 309 | 0 | 533 | 0.11 | 0.03 |  |  | 0.63 | 0.22 |  | 0.20 | 0.08 |  |  | 0.80 | 0.17 |  |
| PRAD | 4 | 50 | 429 | 0.17 | 0.10 | 0.25 | 0.20 | 0.63 | 0.19 | 0.38 | 0.38 | 0.13 | 0.51 | 0.22 | 0.77 | 0.12 | 0.26 |
| HNSC | 56 | 50 | 530 | 0.05 | 0.03 | 0.12 | 0.09 | 0.62 | 0.16 | 0.51 | 0.06 | 0.03 | 0.22 | 0.14 | 0.69 | 0.13 | 0.46 |
| BRCA | 4 | 98 | 748 | 0.05 | 0.01 | 0.21 | 0.16 | 0.62 | 0.21 | 0.40 | 0.03 | 0.01 | 0.38 | 0.21 | 0.73 | 0.16 | 0.35 |
| LUSC | 257 | 43 | 361 | 0.09 | 0.06 | 0.04 | 0.02 | 0.61 | 0.19 | 0.57 | 0.14 | 0.09 | 0.10 | 0.05 | 0.67 | 0.18 | 0.58 |
| LUAD | 257 | 32 | 466 | 0.09 | 0.06 | 0.04 | 0.01 | 0.58 | 0.18 | 0.54 | 0.14 | 0.09 | 0.08 | 0.04 | 0.68 | 0.14 | 0.59 |
| KIRP | 5 | 45 | 226 | 0.02 | 0.01 | 0.16 | 0.08 | 0.57 | 0.24 | 0.40 | 0.01 | 0.01 | 0.47 | 0.12 | 0.73 | 0.18 | 0.26 |
| STAD | 6 | 2 | 339 | 0.05 | 0.07 | 0.38 |  | 0.55 | 0.23 | 0.17 | 0.06 | 0.11 | 0.49 |  | 0.61 | 0.22 | 0.13 |
| DLBC | 0 | 0 | 48 |  |  |  |  | 0.51 | 0.29 |  |  |  |  |  | 0.56 | 0.31 |  |
| GBM | 309 | 2 | 142 | 0.11 | 0.03 | 0.06 |  | 0.49 | 0.27 | 0.44 | 0.20 | 0.08 | 0.36 |  | 0.69 | 0.23 | 0.32 |
| MESO | 0 | 0 | 87 |  |  |  |  | 0.47 | 0.24 |  |  |  |  |  | 0.61 | 0.21 |  |
| SKCM | 2 | 2 | 462 | 0.07 | 0.02 | 0.25 |  | 0.44 | 0.30 | 0.20 | 0.13 | 0.05 | 0.46 |  | 0.55 | 0.32 | 0.09 |
| SARC | 2 | 4 | 245 | 0.07 | 0.00 | 0.13 | 0.11 | 0.44 | 0.33 | 0.31 | 0.35 | 0.05 | 0.24 | 0.18 | 0.59 | 0.33 | 0.35 |
| PAAD | 4 | 10 | 146 | 0.25 | 0.15 | 0.24 | 0.19 | 0.39 | 0.21 | 0.15 | 0.58 | 0.20 | 0.47 | 0.23 | 0.50 | 0.21 | 0.03 |
| KIRC | 5 | 160 | 325 | 0.02 | 0.01 | 0.16 | 0.07 | 0.35 | 0.23 | 0.19 | 0.01 | 0.01 | 0.44 | 0.13 | 0.53 | 0.23 | 0.09 |
| COAD | 19 | 38 | 303 | 0.14 | 0.07 | 0.04 | 0.01 | 0.34 | 0.32 | 0.30 | 0.18 | 0.09 | 0.04 | 0.03 | 0.38 | 0.33 | 0.34 |
| LAML | 28 | 0 | 194 | 0.04 | 0.02 |  |  | 0.33 | 0.36 |  | 0.05 | 0.04 |  |  | 0.38 | 0.38 |  |
| TGCT | 4 | 0 | 156 | 0.05 | 0.02 |  |  | 0.32 | 0.29 |  | 0.02 | 0.01 |  |  | 0.36 | 0.32 |  |
| ACC | 5 | 0 | 80 | 0.03 | 0.02 |  |  | 0.28 | 0.32 |  | 0.02 | 0.01 |  |  | 0.40 | 0.38 |  |
| READ | 0 | 7 | 99 |  |  | 0.05 | 0.02 | 0.26 | 0.30 | 0.21 |  |  | 0.04 | 0.01 | 0.31 | 0.33 | 0.27 |
| KICH | 5 | 0 | 66 | 0.02 | 0.01 |  |  | 0.20 | 0.22 |  | 0.01 | 0.01 |  |  | 0.40 | 0.31 |  |
| UVM | 0 | 0 | 80 |  |  |  |  | 0.16 | 0.20 |  |  |  |  |  | 0.31 | 0.31 |  |
| THCA | 0 | 56 | 515 |  |  | 0.04 | 0.03 | 0.11 | 0.15 | 0.06 |  |  | 0.10 | 0.09 | 0.24 | 0.25 | 0.14 |
| THYM | 2 | 2 | 124 | 0.02 | 0.00 | 0.04 |  | 0.09 | 0.16 | 0.05 | 0.02 | 0.00 | 0.04 |  | 0.12 | 0.21 | 0.09 |
| PCPG | 0 | 3 | 184 |  |  | 0.03 |  | 0.06 | 0.12 | 0.03 |  |  | 0.02 |  | 0.12 | 0.23 | 0.09 |

Since the nonmalignant tissues from TCGA demonstrated reduced but significant SCT promoter methylation (Fig. 3 and Fig. S6 middle panel), we collected DNA methylation data from cancer free samples (Table S2) to account for possible field effect. As can be seen from Fig. 3 and Fig. S6 bottom panel, it appears that SCT promoter is not methylated in corresponding normal tissues/organs, though the sample sizes for a few tissue types are small. Moreover, SCT promoter does not appear to be methylated in 98% samples from >30 tissue/organ types from cancer free samples (Table S2; Fig. S7).

Based on the above data, we concluded that SCT promoter methylation is a highly discriminative biomarker for lung and fourteen other cancers. Interestingly, SCT gene does not appear to be expressed in the primary tumor and non-malignant samples in our analysis (Table S6), In turn, we did not detect SCT differential expression between tumor and non-malignant samples in our analysis (Fig. S8).

## III. DISCUSSION

In this report, we have discovered that SCT promoter methylation is a potential highly discriminative biomarker to separate tumors from non-malignant tissues for lung and fourteen other malignancies. Given the great discriminative potential of SCT promoter methylation in these fifteen cancers, applications for non-invasive procedures such as detection in urine for bladder or renal cancers, sputum (or nasal mucosa as a surrogate marker) for lung cancer [19][20], nipple aspirates for breast cancer, prostatic massage for prostate cancer may also be feasible.

## IV. MATERIALS AND METHODS

### A. Cell lines and DNA

The 15 NSCLC lung cancer cell lines (Table S4) and 3 CDK4/hTERT-immortalized human respiratory epithelial cell lines (Table S4; [21]) were provided from Adi F Gazdar and John D Minna (UT Southwestern Medical Center at Dallas) and the authenticity of these cell lines was confirmed by DNA fingerprint genotyping tests. Genomic DNA was isolated using QIAamp Blood Mini kit (Qiagen). DNA concentrations were measured using Nanodrop2000 (Thermo Scientific).

### B. SCT-specific qMSP

The SCT-specific qMSP assay was designed to target the SCT promoter region and 1st exon and was performed similarly as described previously [22]. The primers and probes sequences are available upon request. The qMSP PCR assay, including MYOD1 gene as input DNA reference [22], was carried out by LightCycler 480 system (Roche). CpGenome Universal Methylated DNA (Millipore) was used as positive control and methylated reference. The level of SCT methylation of a given test was presented as percent of methylated reference (PMR) as determined by the method as described similarly before [22][23]. The methylation status of SCT in these lung cancer cell lines and HREC lines was also confirmed by Illumina HumanMethylation450 BeadChip (Illumina, San Diego, CA).

